# Discovery of a role for *Rab3b* in habituation and cocaine induced locomotor activation in mice using heterogeneous functional genomic analysis

**DOI:** 10.1101/2020.04.21.048405

**Authors:** Jason A. Bubier, Vivek M. Philip, Price E. Dickson, Guy Mittleman, Elissa J. Chesler

## Abstract

Substance use disorders are prevalent and present a tremendous societal cost but the mechanisms underlying addiction behavior are poorly understood and few biological treatments exist. One strategy to identify novel molecular mechanisms of addiction is through functional genomic experimentation. However, results from individual experiments are often noisy. To address this problem, the convergent analysis of multiple genomic experiments can prioritize signal from these studies. In the present study, we examine genetic loci identified in the recombinant inbred (BXD RI) genetic reference population that modulate the locomotor response to cocaine. We then applied the GeneWeaver software system for heterogeneous functional genomic analysis to integrate and aggregate multiple studies of addiction genomics, resulting in the identification of *Rab3b*, as a functional correlate of the locomotor response to cocaine in rodents. This gene encodes a member of the RAB family of Ras-like GTPases known to be involved in trafficking of secretory and endocytic vesicles in eukaryotic cells. The convergent evidence for a role of *Rab3b* was included co-occurrence in previously published genetic mapping studies of cocaine related behaviors; methamphetamine response and *Cartpt* (*Cocaine- and amphetamine-regulated transcript prepropeptide)* abundance; evidence related to other addictive substances; density of polymorphisms; and its expression pattern in reward pathways. To evaluate this finding, we examined the effect of RAB3 complex perturbation in cocaine response. B6;129-*Rab3b*^*tm1Sud*^ *Rab3c*^*tm1sud*^ *Rab3d*^*tm1sud*^ triple null mice *(Rab3bcd*^*-/-*^) exhibited significant deficits in habituation, and increased acute and repeated cocaine responses. This previously unidentified mechanism of the behavioral predisposition and response to cocaine is an example of many that can be identified and validated using aggregate genomic studies.

Many genetic and genomic studies have been performed over the past few decades, representing a wealth of data on the underlying neurobiological and genetic basis of multiple complex behaviors. However, these studies, particularly legacy studies using older technologies and resources lack precision. By aggregating multiple studies, convergent evidence for shared molecular mechanisms of multiple behaviors can be found, for example the widely reported relations among psychostimulant use and novelty response behavior. Here a legacy genetic mapping result for a cocaine related trait mapped in mice was refined using data from 113 different experimental gene sets related to addiction in the GeneWeaver system for heterogeneous functional genomic analysis. Convergent evidence revealed a role for *Rab3b* in this and other traits including multiple psychostimulant responses and CART expression. Experimental perturbation of the RAB complex revealed effects on habituation to a novel environment, cocaine induced activation and *Carpt* expression. The analysis of aggregate data thus revealed a molecular mechanism that influences the relationship between response to novel situations and cocaine-related phenotypes.

## INTRODUCTION

Addiction presents a substantial threat to public health, with 15.3 million persons world-wide experiencing a drug use disorder [1]. Substance use disorders are complex behavioral processes with varying but largely unknown molecular etiology. The self-initiated first use of drugs and progression to addiction are distinct traits that are highly heritable [2]. Such heritable traits can be exploited to identify new biological mechanisms underlying the complex processes that determine addiction behavior, which, in turn, may reveal biomarkers of addiction and/or therapeutic targets for treatment. However, identifying and characterizing the specific effects of genes and variants associated with addiction behavior has been challenging due to the difficulty of modeling addiction in experimental systems [3], and the significant power requirements necessary for human genetic analysis. Genetic and genomic strategies are promising but often yield noisy data with numerous false positives and false negative results. To overcome these barriers, complex, diverse data sets can be aggregated and analyzed using advanced systems genetics approaches to discover the biological mechanisms that are associated with addiction across experimental contexts.

The locomotor activating and sensitizing effect of psycho-stimulants such as cocaine is a well-established behavioral assay for acute drug response, often assayed using the open field device. Open field testing of locomotion has been effective in high-throughput mouse phenotyping experiments to map and identify biological mechanisms underlying response to novel environments and cocaine behavioral responses [4; 5]. Studies in humans and non-human primates have established a relationship between cocaine sensitization with clinical symptomology [6]. Cocaine-induced behavioral sensitization is a measure of drug-induced plasticity, specifically adaptation of the mesocorticolimbic dopamine (DA) system [7]. Locomotion as a behavioral assay of cocaine use was employed to identify *Cyfip2* [8] via QTL mapping in a reduced complexity cross, as a causative variant for acute and sensitized cocaine response phenotypes. Through the use of a systems genetic analysis of cocaine self-administration in mice, *Cyfip2* has recently been associated with a homolog of a psychostimulant addiction candidate, *Fam53b* [9], the ortholog of which was initially identified in a human cocaine dependence GWAS study [10], suggesting that a common and conserved biological mechanism supports sensitization and drug use. These findings indicate that locomotor activation and sensitization to cocaine is an effective phenotypic construct to identify conserved functional genetic correlates of addiction-related behavior.

These assays have been deployed in numerous genetic studies of psychostimulant and other drugs, but efforts to identify trait relevant genes and mechanisms in model organisms has been a lengthy process. The widespread utilization of QTL mapping and functional genomics has resulted in large numbers of QTLs, harboring numerous candidate genes, but for many of these studies, particularly hundreds of legacy studies, the challenging task of identifying the causative genes remains. In a large study using the C57BL/6J x DBA/2J recombinant inbred (BXD RI) genetic reference population and phenotype data from over 250 measures related to multiple behavioral assays across multiple batteries, Philip et al (2010) identified two QTLs (Chr 4 and Chr 15) related to locomotor activation and sensitization. Typical of many genetic analyses in two-progenitor mapping panels these genetic loci are large and require refinement. The interval of the previously reported QTL was 20 Mbp and contained numerous protein-coding genes.

Heterogeneous functional genomic experiments can be integrated to prioritize the many genes in the interval based on diverse evidence sources. GeneWeaver (www.geneweaver.org), facilitates convergent analyses across multiple genomic experiments, platforms and species [11; 12]. The GeneWeaver platform objectively integrates diverse experimental outcomes *in silico* through advanced statistical techniques to provide plausible evidence for previously unknown roles for genes involved in addiction-related behaviors. Genomic databases include published and user submitted experimental data as well as data from multiple large-scale public data resources. We have previously used GeneWeaver to identify a gene, *Ap3m2*, underlying two overlapping biological phenomena, alcohol withdrawal and alcohol preference [13], that we have validated using *Ap3m2* null mice. These and other studies (for review, Bubier et al, 2015 [14]) demonstrate the effectiveness of this system to find a convergent signal in noisy functional genomics data.

In the present study, we sought to refine the large QTLs previously detected for cocaine-induced locomotor activation [15] using refined QTL analysis followed by gene prioritization using heterogeneous functional genomic studies. We prioritized positional candidates using convergent evidence from a database of publicly available genotypes, expression data, sequence variation, ontology annotations and genome wide experiments from the addiction genomics literature to nominate the most likely candidate(s) in an interval for experimental evaluation. We identified a promising mechanistic candidate gene, *Rab3b*, and evaluated its role in cocaine-induced locomotor activation and sensitization using a gene knockout approach.

## MATERIALS AND METHODS

### QTL Mapping in the BXD RI

Quantitative trait locus (QTL) mapping of the previously identified acute locomotor response to cocaine [15] was performed in combined male and female mice from the BXD recombinant inbred strain panel using GeneNetwork (RRID:SCR_002388) [15; 16; 17]. The original analysis of these data was based on an earlier marker map and performed a single-locus scan. To confirm the original result, and more precisely model the effect and locations of variation influencing this trait, R/QTL (RRID:SCR_009085) was utilized for additional QTL model fitting [18] of multiple loci. One thousand permutations were performed for each mapping analysis to obtain genome-wide suggestive and significant thresholds. Ninety-five percent confidence intervals (CI) were obtained by extracting 1.5-LOD drop around the peak marker for statistically suggestive and/or significant QTLs.

### Integrative functional genomic analysis of the candidate genes using GeneWeaver

#### Database

The candidate genes residing within the 1.5 LOD drop interval of detected QTLs were imported into GeneWeaver, (RRID:SCR_003009). At the time of the analysis, the GeneWeaver database contained ∼ 100,000 gene sets. The database consists of diverse curated gene sets. Mammalian phenotype (MPO, RRID:SCR_004855) and gene ontology (GO, RRID:SCR_006447) annotations and previously published QTLs were obtained from the Mouse Genome Database (MGD, RRID:SCR_012953). Published gene expression studies were obtained from either the Neuroscience Information Framework [19] (RRID:SCR_00483) directly from publications. Basal gene expression from striatal and nucleus accumbens obtained from Allen Brain Atlas (RRID:SCR_005984), polymorphic genes among the BXD RI reference panel, and, BXD gene expression correlates for striatal and nucleus accumbens to locomotor response after the first and second *ip* injection of 10 mg/kg of cocaine were generated using GeneNetwork. To find convergence of experimentally derived gene associations from genomewide experiments, the database was queried for “Cocaine, Nicotine or Methamphetamine” followed by manual review to omit false positive search results, e.g., those for which cocaine, nicotine or methamphetamine was mentioned in the publication abstract but was not relevant to the specific gene set. This convergent analysis resulted in the retrieval of 113 data sets.

#### Gene prioritization

GeneWeaver’s GeneSet graph tool represents the collections of gene lists as a bipartite graph, from which high-degree vertices can be identified. This structure is analyzed using the Hierarchical Similarity (HiSim) graph tool to enumerate all intersections among gene sets and highlight high order intersections containing positional candidates for prioritization of candidate genes within the two chromosomal intervals. This was performed by interrogating each of the two candidate gene lists, one from each locus, with selected gene sets describe above. HiSim was used to group experimentally derived gene-set results based on the genes they contain. For a collection of input gene sets, this tool constructs a graph of hierarchical relationships in which each terminal node represents individual gene sets and each parent node represents gene-gene set bicliques found among combinations of these sets using the maximal biclique enumeration algorithm (MBEA) [20]. The resulting graph structure was determined solely from the gene-set intersections of every populated combination of gene sets. In terms of gene sets, the smallest intersections (fewest gene sets, most genes) were at the lower levels, and the largest intersections (most gene sets, fewest genes) were at the top of the graph. To prune the hierarchical similarity graph, bootstrapping was performed. The graph in the present analysis was sampled with replacement at 75% for 1,000 iterations; node-node parent-child relationships occurring in greater than 50% of the results were included in the bootstrapped graph.

### Animals

Generation of mice with targeted deletions in *Rab3a; Rab3b; Rab3c* and/or *Rab3d* function has been previously described [21]. *Rab3a* heterozygous and *Rab3bcd* triple null mice were obtained from The Jackson Laboratory (Strain Name: B6;129-*Rab3b*^*tm1Sud*^ *Rab3a*^*tm1Sud*^ *Rab3d*^*tm1Rja*^ *Rab3c*^*tm1Sud*^/J; RRID:IMSR_JAX:006375) and bred to produce obligatory *Rab3bcd* triple heterozygous (*Rab3bcd*^+/-^) and triple null (*Rab3bcd*^-/-^) offspring. Ninety-six mice belonging to three genotype groups, both sexes and two treatments (saline or cocaine) were tested. Age- and sex-matched mice were randomly assigned to the following groups: C57BL/6J (controls RRID:IMSR_JAX:000664); *Rab3bcd*^*+/-*^; and *Rab3bcd*^*-/-*^. Mice were between 76 and 116 days of age at the start of behavioral testing. All mice were maintained at The Jackson Laboratory in climate- and 12 h light cycle-controlled rooms and provided acidified water and 5K52 chow (LabDiet/PMI Nutrition, St. Louis, MO) *ad libitum*. The Jackson Laboratory Animal Care and Use Committee approved all protocols involving mice.

### Behavioral Analysis

Prior to testing, mice were randomly assigned either to a saline (SAL) or cocaine (COCA) group. Mice were brought from their home room into the test room and allowed to habituate for one hour. Mice were administered 10 mg/kg of saline or cocaine by *ip* injection. Immediately post injection, mice were placed in an open field box (39 × 39 × 39 cm) and locomotor activity was recorded over 20 minutes using the EthoVision XT 8 (RRID:SCR_000441) video tracking and analysis system (Noldus Information Technology, Wageningen, The Netherlands). On days 1 and 2 of testing all mice received saline. Day 1 measures locomotor activity in response to injection stress and novel environment, while Day 2 measures locomotor activity within a familiar environment and attenuated injection stress. On days 3, 5, 7 and 9, mice received their assigned saline or cocaine injection and the development of cocaine sensitization was measured relative to activity observed on Day 3 Tester was blind to all genotypes and treatments. Data were binned into intervals of five minutes each.

### Tissue Collection

Whole striatal samples were collected to evaluate expression of *Cartpt*. In order to minimize differences in gene expression due to stress from handling and injection, mice were sacrificed via decapitation within 24 hrs of the last behavioral trial. In addition to mice that were subjected to cocaine and saline treated mice from behavioral testing, a behaviorally and drug naïve group of mice (n=8) were also sacrificed via decapitation to obtain basal levels of C*artpt* and housekeeping genes. Striatal regions were dissected under a dissecting scope and total striatal RNA was isolated.

### qRT-PCR

Striatal tissue samples were stored in RNAlater (Life Technologies, Carlsbad, CA), homogenized and total RNA was isolated by standard TRIzol® Plus (Life Technologies, Carlsbad, CA) methods that included DNase digestion. The quality of the isolated RNA was assessed using an Agilent 2100 Bioanalyzer instrument (RRID:SCR_018043) and RNA 6000 Nano LabChip assay. 500ng of total RNA was then reverse transcribed with random decamers and M-MLV reverse transcriptase (RT) using the Message Sensor RT Kit (Life Technologies, Carlsbad, CA).

### Statistical analysis

Statistical analyses were performed in JMP 9.0 (RRID:SCR_014242 SAS Institute Inc., Raleigh, NC). To analyze the locomotor data, we performed a 3 × 2 × 2 × 6 mixed-model ANOVA using distance traveled over 20 min as the dependent measure. Between-subjects factors were *Rab3bcd* genotype (+/+, +/-, -/-), drug (saline, cocaine), and sex. Session (1-6) was a within-subjects factor. To dissociate the effects of *Rab3bcd* complex deletion on habituation and cocaine sensitization we performed a 3 × 2 × 2 × 6 mixed-model ANCOVA using distance traveled on session 1 and session 2 as covariates and using the same dependent measure and independent factors used in the initial ANOVA.

## RESULTS

### QTL Mapping

Locomotor activity in response to cocaine (*ip* injection of 10 mg/kg of cocaine) was genetically mapped to loci on Chr 4 and 15 in a previous study [15]. The data from this study was remapped with software allowing for the inclusion of both effects in the same model, ensuring improved accuracy of the QTL locations. The overall QTL model that best predicted locomotor response following repeated injection of cocaine (GeneNetwork Dataset GN11796; Fig. 1A) was a female-specific two-QTL additive model involving Chr 4 (*Cocia15 Cocaine Induced Activation 15*, rs13477919 @ 104.96 Mb) and Chr 15 (*Cocia16 Cocaine Induced Activation 16* rs13482528 @ 38.82 Mb), which together accounted for 34.28 % of the trait variance (Fig. 1B). A 1.5 LOD window was used to define the 95% confidence interval (CI). The Chr 4 CI spans 101.10 (rs13477873) – 114.51 (gnf04.110.583) Mb while the Chr 15 CI spans 33.43 (rs8267966) – 45.11 (rs13482547) Mb. The Chr 4 and Chr 15 QTLs accounted for 11.38 % and 15.60 % of the trait variance, respectively. No significant interactions among these loci were detected. Positional candidate genes underlying each of these regions were retrieved from the Mouse Genome Informatics (MGI) database (http://informatics.jax.org) [22]. Each gene list was uploaded to GeneWeaver.org [12] for integrative functional genomic analysis.

**Figure 1:**
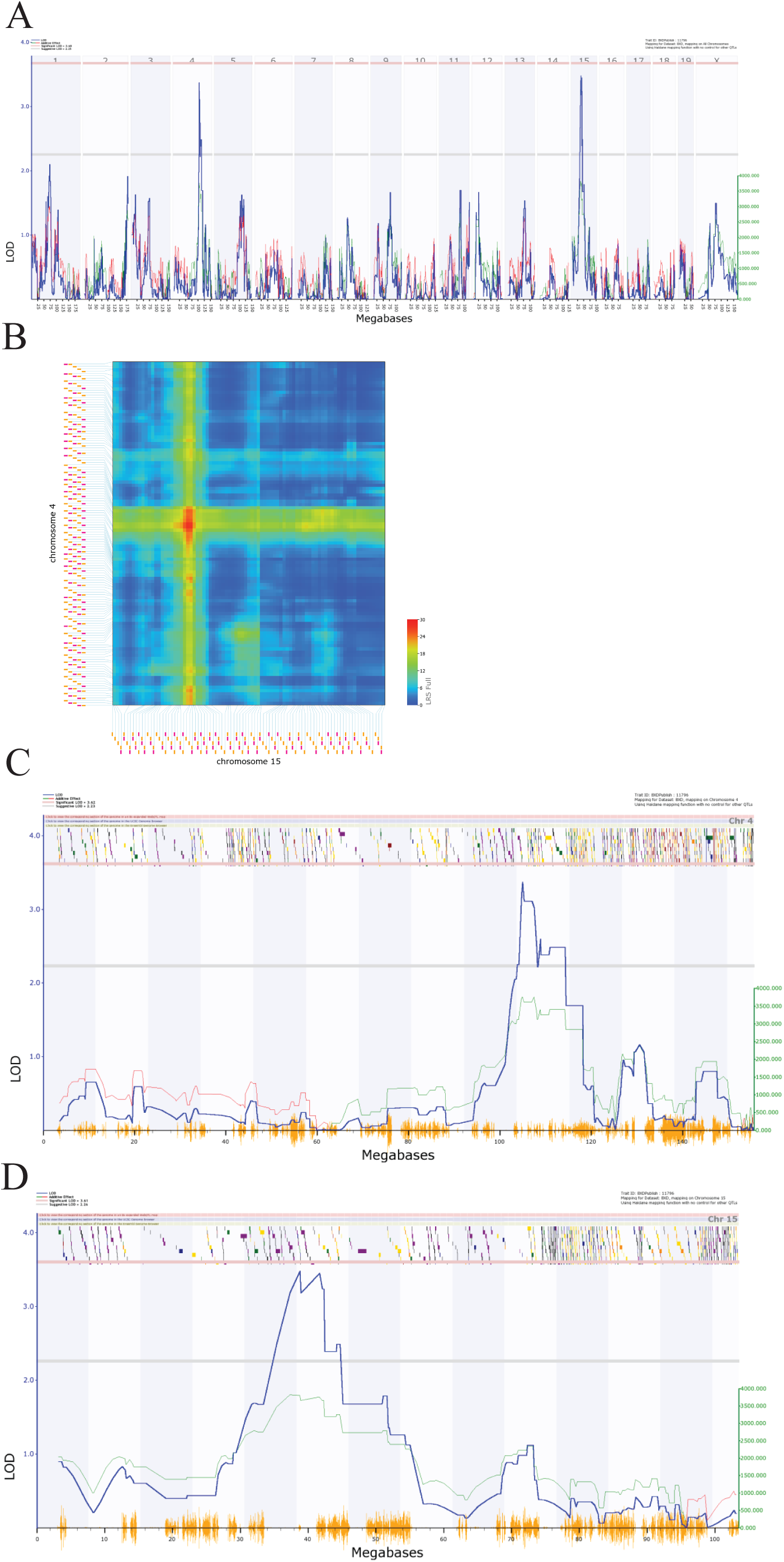
QTL mapping of cocaine response in the expanded BXD RI genetic reference population: QTL mapping of locomotor response following the second injection of cocaine in females resulted in the detection of suggestive QTLs on Chr 4 and Chr 15. A). Green lines indicate regions where DBA/2J is associated with increased phenotypic effect the increaser allele, red lines indicate that C57BL/6J is associated with the increased phenotypic effect. B) the two QTL additive model involving Chr 4 and Chr 15. Additive the black horizontal lines represent the 1.5 LOD support interval on Chr 4 (C) and Chr 15 (D). The LOD support interval on Chr 4 spans 101.10 (rs13477873) 114.51 (gnf04.110.583) Mb, while on Chr 15 it spans 33.43 (rs8267966) 45.11(rs13482547) Mb. 108 and 61 positional protein coding candidate genes reside in the Chr 4 and Chr 15 intervals, respectively.

### Integrative functional genomic analysis of the candidate genes reveals *Rab3b* as a likely candidate gene for locomotor response to cocaine

A search of keywords pertaining to cocaine, methamphetamine and nicotine was performed in GeneWeaver [05-01-2012] and 3,315 sets were retrieved. This was followed by a closer inspection of the descriptive meta-content associated with the search results. This was done to remove redundant gene sets and misidentified gene sets (e.g. abstract referenced cocaine, gene set was from untreated controls). A total of 113 gene sets with an average of 66.5 genes per gene set and spanning 19 publications (Fig. 2A, Table S1) were retained for analysis. The HiSim analysis tool was used to construct an empirically derived hierarchy of the input gene sets for both the Chr 4 and Chr 15 candidate gene list. This allowed us to identify the most highly connected positional candidate genes among a large number of genomic data sets, suggesting that it is the one most frequently implicated in functionally relevant studies. The gene with the highest degree (7) of connectivity from either the Chr 4 (Fig. 2B) and Chr 15 positional candidates was *Rab3b*, a member of the RAS oncogene family thought to function in protein metabolism and cell junction dynamics[23; 24]. *Rab3b* is expressed in the dorsal and ventral striatum, down-regulated in C57BL/6J and up-regulated in C3H/HeJ after exposure to another psychostimulant, nicotine GS:14888 [25], and is differentially expressed in dopamine receptor 1 mutants when compared to their wild-type controls after saline treatment GS:87050 [26]. Furthermore, *Rab3b* is a candidate gene for three previously published QTLs, all of which were mapped in the BXD RI reference population, derived from the inbred progeny of a cross of the C57BL/6J and DBA/2 strains. These traits were climbing behaviors in response to methamphetamine GS:84160 [27], repeated movements following cocaine GS:84158 [28] and hypothalamic CART peptide (*Carpt*) abundance GS:83998 [29]. In each of the three QTLs, the DBA/2 derived allele increases activity and peptitde release, which in the same direction as the allelic effect on cocaine response QTL on Chr 4 (*Cocia15*).

**Figure 2:**
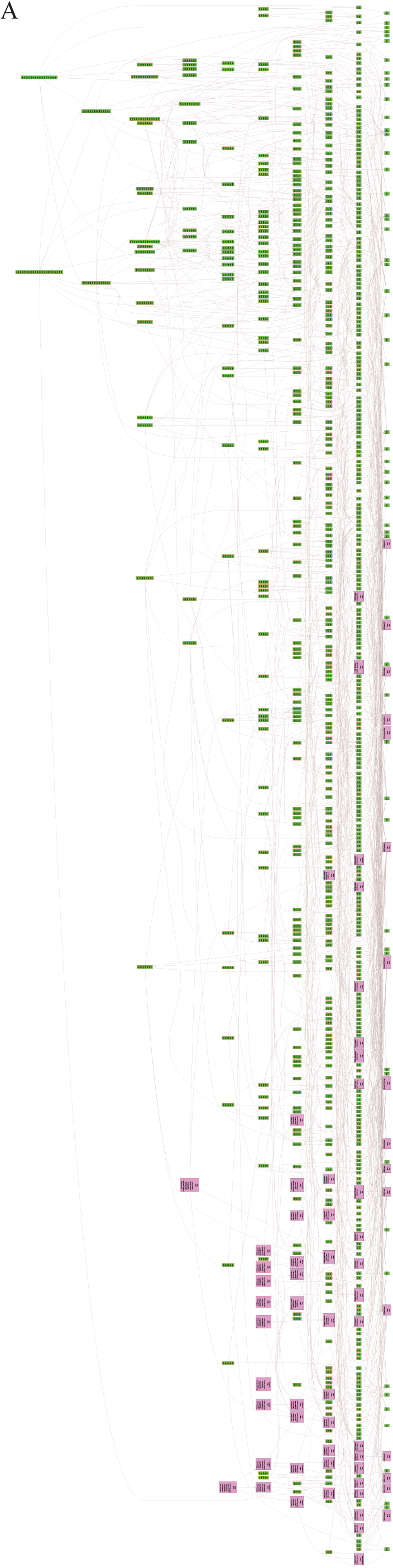
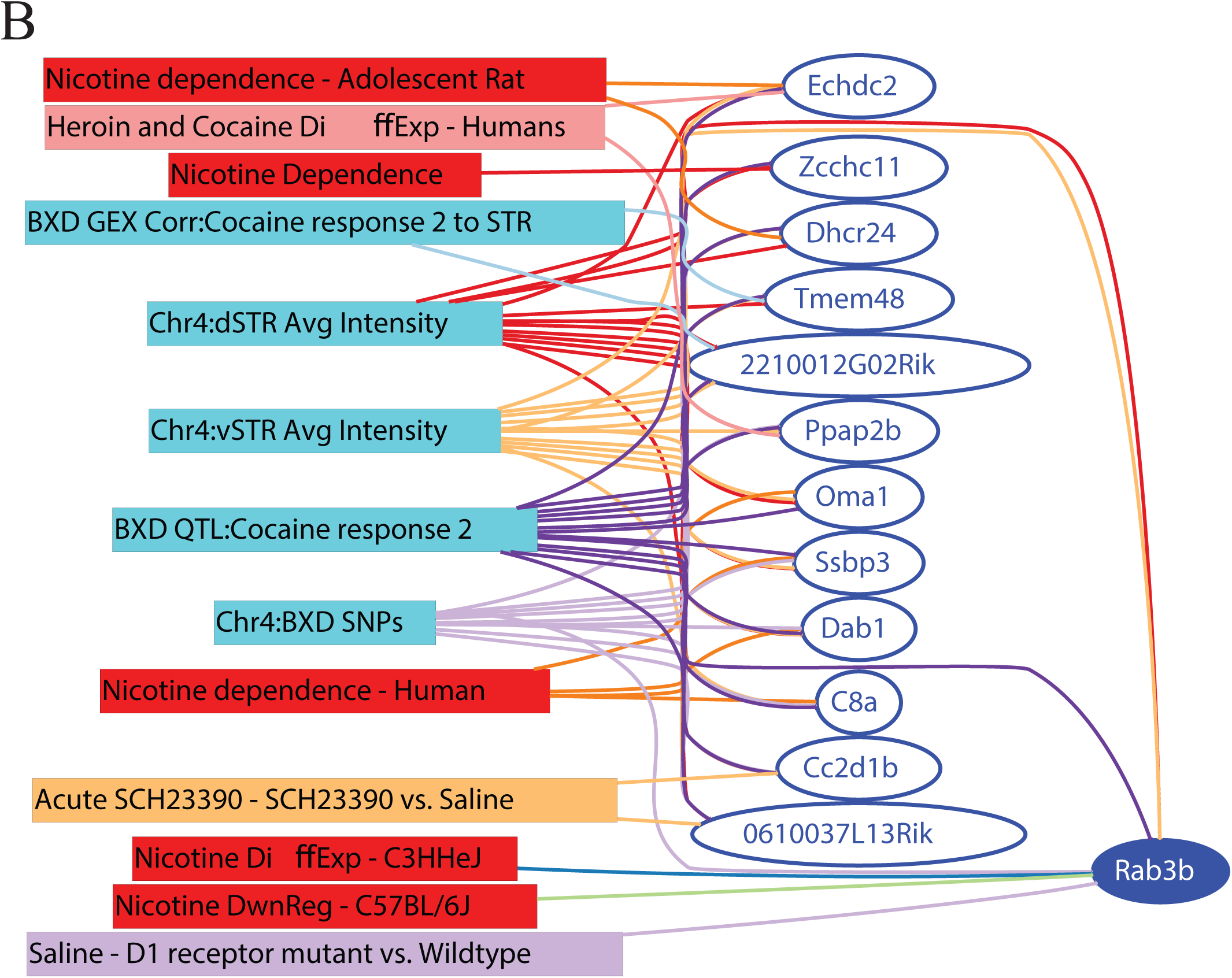
Integrative functional analysis in GeneWeaver: A) Hierarchical Similarity tool depicting a hierarchical representation of gene sets used in the integrative analysis. Nodes at the top are highly connected genes. Pink colored nodes represent nodes containing one or more QTL candidate genes. Rab3b is revealed as the most highly connected QTL candidate gene. B) GeneSet graph representing the underlying interconnections among the gene sets containing Rab3b.

To evaluate this candidate experimentally, *Rab3b* targeted mice were used. Previous studies reveal that compensatory effects of Rab complex proteins limit the effect of individual gene deletions, therefore a triple null mouse strain involving *Rab3b, Rab3c* and *Rab3d* B6;129-*Rab3b*^*tm1Sud*^ *Rab3c*^*tm1sud*^ *Rab3d*^*tm1sud*^ [21] was obtained from The Jackson Laboratory for experiments.

### Assessment of Cocaine Response in *Rab3bcd* Triple Null mice

A nine day cocaine sensitization paradigm modeled after the work of Phillips et al. [30] was administered. Briefly, on days 1 and 2 of testing all mice received saline. Day 1 measures locomotor activity in response to injection stress and novel environment, while Day 2 measures locomotor activity within a familiar environment (habituation) and attenuated injection stress. On Days 3, 5, 7 and 9, mice received their assigned saline or cocaine injection *i*.*p*., blinded to investigator.

There were significant main effects of genotype [F_(2, 83)_ = 10.68, p < 0.0001], drug [F_(1, 83)_ = 194.69, p < 0.0001], and session [F_(5, 79)_ = 101.31, p < 0.0001], as well as a significant 3-way interaction of these factors [F_(10, 158)_ = 1.98, p = 0.03]. To determine the nature of the 3-way interaction, we examined performance of each of the genotype-drug subgroups on all sessions (Fig. 3A, B, C). *Rab3bcd*-/- mutants exhibited significantly greater cocaine activation (p <0.05) relative to wild type and heterozygote mice on all but the second cocaine session (Fig. 3B). However, *Rab3bcd*-/- mutants also traveled a significantly greater distance relative to wild type and heterozygote mice on several sessions in the saline condition beginning with the first saline injection (Fig. 3C), suggesting an effect of *Rab3bcd* complex perturbation on habituation to a novel environment.

**Figure 3.**
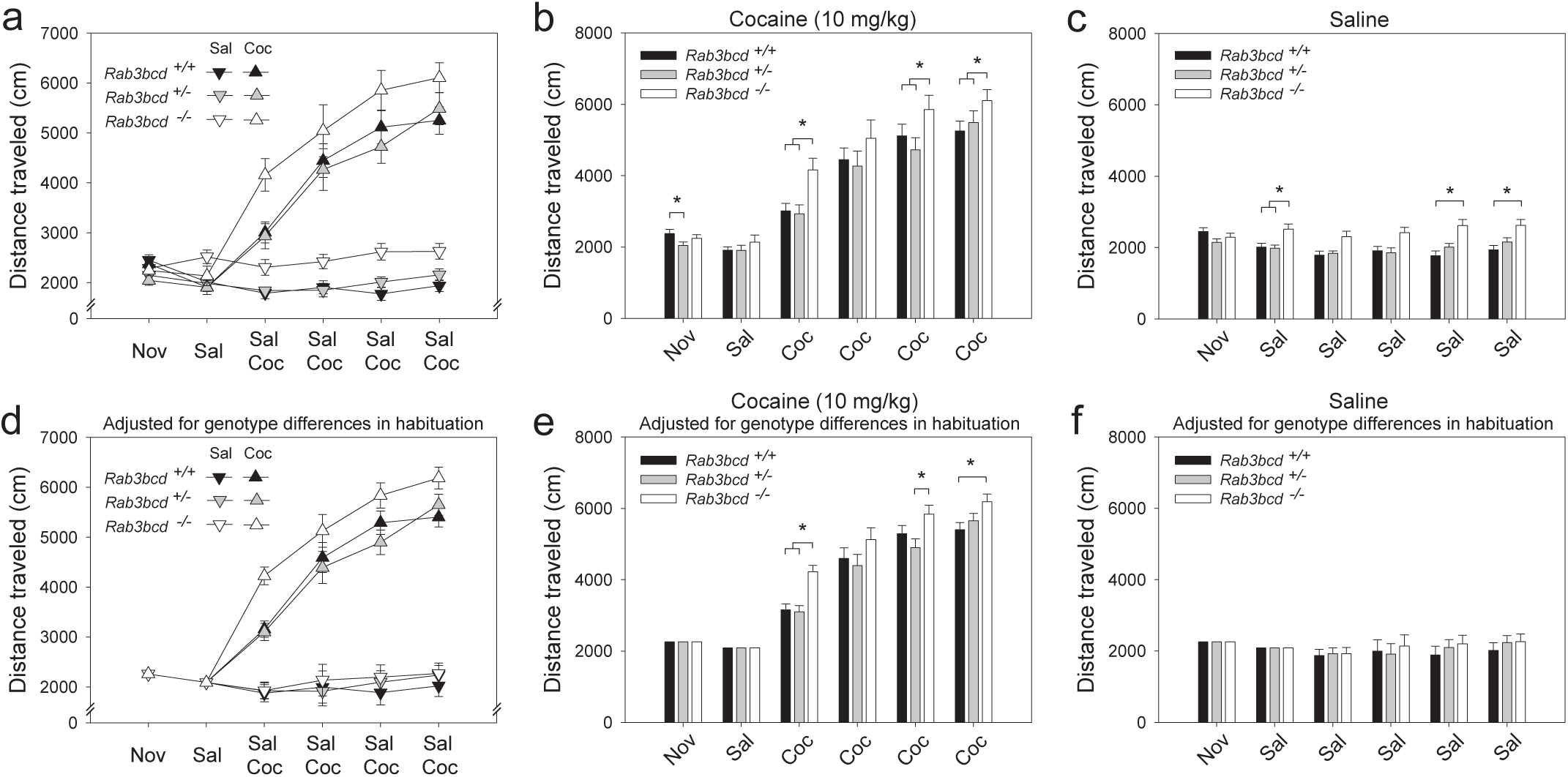
Cocaine induced locomotor activation in Rab3bcd mutants, heterozygotes, and wildtypes. Rab3bcd-/- mutants exhibited significantly greater cocaine sensitization (p < 0.05) relative to wildtypes and heterozygotes on all but the second cocaine session (b). However, Rab3bcd-/- mutants also traveled a significantly greater distance relative to wildtypes and heterozygotes on several sessions in the saline condition beginning with the first saline injection (c), suggesting an effect of Rab3bcd on habituation to the apparatus or injection stress. After adjusting for the effect of habituation (d, e, f) by including distance traveled on the first two sessions as covariates, the significant effect of Rab3bcd on cocaine sensitization persisted, whereas significant differences in distance traveled between wildtypes, heterozygotes, and mutants in the saline condition were no longer observed. These data indicate significant effects of Rab3bcd on both habituation and cocaine sensitization.* p < 0.05

To dissociate the effects of *Rab3b* manipulation on habituation and cocaine sensitization, we performed a 3 × 2 × 2 × 6 mixed-model ANCOVA using distance traveled on session 1 and session 2 as covariates. The main effects of genotype [F_(2, 81)_ = 3.69, p = 0.02], drug [F_(1, 81)_ = 355.83, p <0.0001], and session [F_(4, 78)_ = 6.04, p = 0.0003], as well as the 3-way interaction between these factors [F_(8, 156)_ = 2.24, p = 0.02] remained significant (Fig. 3D). Post-hoc tests indicated that after adjusting for the effects of habituation, effects of *Rab3b* on cocaine activation persisted (Fig. 3E) whereas the previously significant differences in distance traveled between wild types, heterozygote, and mutant mice were no longer observed (Fig. 3F).

### Cartpt transcript abundance in Rab3bcd ^-/-^ and C57BL/6J

The GeneWeaver search revealed a previously published QTL mapping study involving *Cartpt* transcript abundance [29] and the *Cocia15* QTL. *Cartpt* or cocaine- and amphetamine-regulated transcript peptide is involved in reward- and feeding-related behaviors. Studies in rats have shown that CART mRNA expression is upregulated after acute cocaine self-administration in brain regions of the mesocorticolimbic dopamine system [31]. To assess the effects of *Rab3bcd* variation on *Cartpt* expression, we analyzed *Cartpt* expression using ΔC_t_ values to detect genotype (*Rab3bcd* ^*-/-*^ and C57BL/6J) × treatment (naïve, SAL and COCA) effects. The genotype x treatment effect was suggestive (F_(2,2)=_ 2.58; p_genotype × treatment_ < 0.08). No significant differences were observed across genotypes for the naïve group, suggesting that at a baseline line level *Rab3bcd* mutants have no altered expression of *Cartpt*. Planned contrasts revealed differences in *Cartpt* expression following saline or cocaine treatment. Following saline or cocaine treatment *Cartpt* is differentially expressed across the two genotypes (SAL: F_(1,42)_ = 10.03, p_genotype_ < 0.003; COCA: F_(1,42)_ = 11.10, p_genotype_ < 0.002). Within genotype contrasts failed to detect any significant differences across the two treatments (*Rab3bcd* ^-/-^: F_(1,42)_ = 1.76, p_treatment_ < 0.19; C57BL/6J: F_(1,42)_ = 1.84, p_treatment_ < 0.18). These results suggest that disruption of the Rab3 complex induces dysregulation of *Cartpt* in response to injection of either saline or cocaine (Table 1a,b).

**Table 1.**

| Treatment | Naïve |  | SAL |  | COCA |  |
| --- | --- | --- | --- | --- | --- | --- |
| Genotype | C57BL/6J | *Rab3bcd*<sup>-/-</sup> | C57BL/6J | *Rab3bcd*<sup>-/-</sup> | C57BL/6J | *Rab3bcd*<sup>-/-</sup> |
| $\Delta C_t - \mu \pm SE$ | 10.24 ± 0.19 | 10.12 ± 0.23 | 10.55 ± 0.11 | 9.75 ± 0.15 | 10.22 ± 0.16 | 9.42 ± 0.18 |

B
| Treatment | Naïve |  | Injected (SAL or COCA) |  |
| --- | --- | --- | --- | --- |
| Genotype | C57BL/6J | *Rab3bcd*<sup>-/-</sup> | C57BL/6J | *Rab3bcd*<sup>-/-</sup> |
| $\Delta C_t - \mu \pm SE$ | 10.24 ± 0.18 | 10.12 ± 0.22 | 10.36 ± 0.11 | 9.58 ± 0.11 |

## DISCUSSION

Here we present the discovery of a role for *RAB3B* complex in habituation and cocaine induced activation using the GeneWeaver integrative functional genomic analysis of the candidate genes for a previously mapped cocaine-induced activation QTL among BXD RI genetic reference population [15]. We have shown that by incorporating multiple independent lines of evidence, from convergent studies it is possible to readily interrogate a QTL interval, to arrive at highly likely candidate genes. In the integrative functional genomic analysis described in this report, we have incorporated empirical information derived from 21 published studies. The approach is relatively unbiased in that the genomewide studies integrated into the analysis evaluate virtually all positional candidates in the QTL interval. Using these genomic data sets, we have identified a highly likely gene, *Rab3b*, which resides in the *Cocia15* QTL. RAB proteins are proteins belonging to the family of Ras-like GTPases. These proteins are primarily involved in trafficking of secretory and endocytic vesicles of eukaryotic cells [32; 33]. Sixty known RAB proteins exist, with the RAB3 subfamily being the most abundant and expressed at variable levels in the brain [23; 24; 34]. The RAB3 protein subfamily consists of four homologous isoforms, namely, RAB3A, 3B, 3C and 3D. RAB3A, 3B and 3C co-localize on presynaptic vesicles, while RAB3D localizes to secretory vesicles in exocrine glands [35; 36; 37]. The RAB3 subfamily is involved in regulated exocytosis and in docking, priming and fusion stages of the synaptic vesicle cycle. Specifically, *in vitro* studies have revealed that RAB3 proteins inhibit Ca^2+^-triggered exocytosis in PC12 cells [37]. Linkage and association evidence across multiple independent studies has suggested variation in the RAB3B subunit as a source of the trait but deletion of this gene is likely developmentally compensated by other subunits due to functional redundancy, with *RAB3A* minimally needed for survival [21; 38]. Because of this functional redundancy among RAB3 members, a triple knock-out approach was necessary to effectively inactivate this functionality. Of these genes only *Rab3b* resides in the QTL interval on Chr 4. Our *Cocia15* QTL for cocaine response harboring *Rab3b*, has also been detected in three previous mapping studies, namely, CART (*Carpt*) transcript abundance [29], methamphetamine-related climbing behavior [27] and cocaine-related behavior (repeated movements at 5 mg/kg) [28]. *Rab3b* is expressed in the striatum, nucleus accumbens and the ventral tegmental area of mice (www.brain-map.org), all of which are addiction-relevant regions of the brain. Each of these brain regions contains dopaminergic neurons and plays a central role in cocaine-mediated behaviors. *In vitro* and *in vivo* studies involving overexpression of *Rab3b* in dopamine neurons of the rat substania nigra (SN) has revealed that overexpression of *Rab3b* increases dopamine content, uptake, number and size of synaptic vesicles, and levels of presynaptic proteins [38]. The role of *Rab3b* in Ca^2+^-triggered exocytosis, localization on synaptic vesicles, expression in addiction-relevant tissues and appearance in multiple independent QTL mapping studies that target the dopaminergic system makes it a highly likely candidate gene underlying the cocaine-induced locomotor response QTL.

The effects of RAB complex manipulation on habituation to the open field, behavioral response to repeated cocaine injection, and the effects on CART expression which are induced by either saline or cocaine injection suggest that the complex is involved in response to salient or novel experiences, and the potential reinforcement of exposure to certain novel experiences. A significant literature supports the hypothesis that exposure to novelty and sensory stimuli are reinforcing across species [reviewed in 39]. Moreover, multiple studies reveal the predictive relationship of novelty seeking and sensation seeking on drug use and effect [9; 40; 41; 42]. The present findings provide causal evidence that the RAB complex is part of a larger network underlying these associations.

The putative mechanism of variation in the mouse can be discerned from the location of the variants affecting *Rab3b*. The SNPs that exist in *Rab3b* between C57BL/6 and DBA/2 founders of the BXD RI strains are either intronic or 5’ and 3’ intergenic. The Sanger resequencing project, using heterogeneous stock QTL mapping results, has shown that functional variants at small effect QTLs (<4%) are most likely to be intergenic, and that functional variants at large effect QTLs (>4%) are most likely to be intronic [43]. *Cocia15* QTL harboring *Rab3b* is a large effect QTL accounting for 11.38% of the trait variation. There is no cis-eQTL for *Rab3b* detected in the BXD RI panels (Genenetwork.org not shown) and although we tested for differential splicing of the Rab3b-L (OTTMUST00000018722) and Rab3b-S (OTTMUST00000018724) transcripts, we detected no alternate splicing between the drug naive founders (data not shown). The absence of a detectable *Rab3b* eQTL does not preclude genetically allelic variation in expression regulation. One potential explanatory SNP that exists in B6 vs D2 is at rs32262045. Importantly this SNP lies within a region of *Rab3b* known to display cocaine-induced epigenetic changes in 5-hydroxymethylcytosine (5hmC) [44]. These types of cocaine-induced DNA methylation changes genomic regions are correlated with changes in local gene expression. Another promising causal variant is SNP rs32532179 which lies within a LINE/L1 (Long interspersed nuclear element) element of 738 bp LINE [45]. Repetitive elements, in particular LINE1 have recently been shown to be subject to activation or epigenetic regulation through H3K9Me3. These SNPs leave open the possibility that an epigenetic mechanism; with complex developmental and spatial regulation underlies the behavioral phenotype. GeneNetwork does capture many brain regions but we know it is not exhaustive in addition the altered differential expression of *Rab3b* during embryogenesis due to B6/D2 allelic variation can produce the observed phenotypic differences. This would be consistent with the known embryonic lethality of the Rab3 quadruple mutant (Rab3a, Rab3b, Rab3c and Rab3d). Engineering of these variants in cellular and mouse experiments can provide conclusive experimental identification of the regulatory variant, and how it responds to repeated experience.

In summary, our findings demonstrate the use of integrative functional genomic as a tool to identify a novel biological mechanism of cocaine induced locomotor activation, habituation and other effects of exposure. Using the integrative functional genomic analysis we identified and validated of *Rab3b* as a candidate gene for cocaine-induced locomotor activation based on genetic and functional evidence. Future studies into this mechanism may elucidate its precise role in the response to novel stimuli. Further application of the integrative functional genomics approach may enable discovery of mechanism of other complex trait variation from aggregate data from current and legacy genetics and genomics.

## FUNDING

Funding for this project comes from NIAAA R01 AA018776 (supported by NIAAA and NIDA), R01 DA 037927, P50 DA39841 to EJC, and U01 DA043809 to JAB. PED was funded by NIDA K99 DA043573 during preparation of this manuscript.

## ACKNOWLEDGEMENTS

The authors would like to thank Dr. Raymond F. Robledo technical assistance or all the brain dissections. The authors would also like to thank Dr. Brynn H. Voy, Dr. Arnold M. Saxton and Dr. Matthew A. Cooper for their helpful comments on this research. Finally, we thank Stephen Krasinski for assistance with this manuscript.

## Notes

### Competing Interest Statement

The authors have declared no competing interest.

